# Integration, exploration, and analysis of high-dimensional single-cell cytometry data using Spectre

**DOI:** 10.1101/2020.10.22.349563

**Authors:** Thomas Myles Ashhurst, Felix Marsh-Wakefield, Givanna Haryono Putri, Alanna Gabrielle Spiteri, Diana Shinko, Mark Norman Read, Adrian Lloyd Smith, Nicholas Jonathan Cole King

## Abstract

As the size and complexity of high-dimensional cytometry data continue to expand, comprehensive, scalable, and methodical computational analysis approaches are essential. Yet, contemporary clustering and dimensionality reduction tools alone are insufficient to analyze or reproduce analyses across large numbers of samples, batches, or experiments. Moreover, approaches that allow for the integration of data across batches or experiments are not well incorporated into computational toolkits to allow for streamlined workflows. Here we present Spectre, an R package that enables comprehensive end-to-end integration and analysis of high-dimensional cytometry data from different batches or experiments. Spectre streamlines the analytical stages of raw data pre-processing, batch alignment, data integration, clustering, dimensionality reduction, visualization and population labelling, as well as quantitative and statistical analysis. Critically, the fundamental data structures used within Spectre, along with the implementation of machine learning classifiers, allow for the scalable analysis of very large high-dimensional datasets, generated by flow cytometry, mass cytometry (CyTOF), or spectral cytometry. Using open and flexible data structures, Spectre can also be used to analyze data generated by single-cell RNA sequencing (scRNAseq) or high-dimensional imaging technologies, such as Imaging Mass Cytometry (IMC). The simple, clear, and modular design of analysis workflows allow these tools to be used by bioinformaticians and laboratory scientists alike. Spectre is available as an R package or Docker container. R code is available on Github (https://github.com/immunedynamics/spectre).

## INTRODUCTION

### High-dimensional analysis tools

High-dimensional (HD) cytometry plays an important role in the study of immunology, infectious diseases, autoimmunity, hematology and cancer biology, elucidating critical mediators of immunity and disease at a single-cell level. Advances in single-cell cytometry systems (including flow, spectral and mass cytometry) have enabled the simultaneous analysis of over 40 proteins [1] in a single panel, resulting in vast and complex datasets. A large portion of the analysis still utilizes manual gating, which is the sequential and often arduous process of identifying cells, based on the expression of one or two cellular markers at a time. While this allows for user-guided examination of the dataset, it is intractable when mapping out the vast variety of cell phenotypes that may be present in HD datasets, and may introduce selective bias through a subjective and sequential bifurcating inclusion/exclusion of markers [2]. This is in part due to the number of gates required to fully parse the dataset. As such, a variety of computational approaches have been adopted by the cytometry community to help analyze HD datasets, including automated gating [3], clustering (such as PhenoGraph [4], FlowSOM [5], X-Shift [6]), dimensionality reduction (such as t-SNE [7, 8], UMAP [9], trajectory inference (such as Wanderlust [1], Wishbone [10]), and automated cell classification [11] [12, 13]. Many of these tools have been brought together into ‘toolboxes’, providing either code- or GUI-based analysis workflows, such as Cytofkit [14], CITRUS [15], CATALYST [16], Cytofworkflow [17], or diffcyt [18]. In parallel, similar tools have also been developed in other fields, such as a wide variety of analysis approaches for single-cell RNA sequencing (scRNAseq; Seurat [19, 20], SingleCellExperiment [21], Monocle [22]). These have been developed to address similar difficulties in analysis, with developments in one field sometimes assisting another, such as the graph-based clustering from Phenograph [4] informing the design of clustering in Seurat [19]. Overall, these tools allow for automated and data-driven data processing that may be performed in an unsupervised (requiring no human supervision) or semi-supervised manner, requiring some human decision-making.

### Limitations of existing algorithms and toolkits

Despite the advantages offered by these computationally-driven approaches, a number of challenges persist, including slow operation speeds, difficulty in scaling to large datasets, and insufficient reproducibility of analysis results across independent experiments. As such, the use of such tools is often limited to relatively small datasets from single experiments. In particular, many scRNAseq specific tools, while suited to processing datasets with a large number of features (markers), do not necessarily scale well to dataset volumes that include tens to hundreds of millions of cells, as the volume of data may exceed the memory capacity of the computer being used for analysis, and/or the processing may carry excessively long run times [23]. In scRNAseq, independent datasets of this volume are not common outside of cell atlas-type projects [24], but cytometry frequently generates datasets of this size. In addition, these data-driven approaches are highly sensitive to the influence of batch effects, a phenomenon in which cells of the same phenotype differ in their signal intensity across multiple experimental batches. Batch effects are an artefact of time and context, in which experimental conditions and/or instrumental performance inadvertently influence the measured signal intensity. Left uncorrected, computational algorithms may falsely identify differences (or similarities) based on batch, rather than biologically relevant (and correct) differences between samples or experimental groups. Hence, having the ability to control for and differentiate batch effects from biological differences is critical. While a variety of approaches exist for scRNAseq data [19, 20, 25–27], approaches designed for cytometry data are less well developed. While there exist several approaches to align data from different batches, many of these align samples individually against each other in the context of standardized clinical profiling [28], potentially removing important biological variations between samples. In contrast, recent approaches calculate batch adjustments using reference controls: aliquots of a control sample run with each batch. The resultant conversions can then be applied to all samples in the batch, preserving biologically relevant differences [29]. Because some batch effects can differentially impact select cell lineages [30], further techniques extend this reference-based approach to apply adjustments in a cluster-specific manner [30]. As this is a developing area, a number of other approaches have also been proposed, using a variety of methods to remove technical variance [11, 31], some inspired by approaches in scRNAseq.

While a number of clustering, dimensionality reduction, and batch alignment approaches exist, many of these are not directly integrated with each other - and many implementations require specific dataset formats, limiting the flexibility and interoperability of these tools. Specifically, tools that are developed in different fields (e.g. cytometry or scRNAseq) are often provided as stand-alone packages that operate on custom data formats (e.g. flowFrames, SingleCellExperiment objects, Seurat objects, etc). The dependency of individual tools on custom data formats also makes it non-trivial to apply computational tools from a broader data science and machine learning field that are not specifically designed for cytometry (and thus have no knowledge of the custom data format) but are of significant advantage in existing analysis pipelines. A streamlined solution that can operate on large datasets at speed and integrate tools across fields in a modular fashion would be of significant advantage to the cytometry community.

### Spectre for analysis of large and complex high-dimensional cytometry datasets

To address these challenges, we developed Spectre, a computational toolkit in R that facilitates rapid and flexible analysis of large and complex cytometry datasets. This toolkit expands on the ‘Cytometry Analysis Pipeline for large and compleX datasets’ (CAPX) workflow that we have published previously for deep profiling of hematopoietic datasets [32]. Through specific function and workflow design, we demonstrate intuitive, modular, and high-speed workflows for data pre-processing (such as scaling/transformation), clustering, dimensionality reduction, plotting, as well as quantitative statistical analyses. Additionally, we incorporate an effective method for integrating data across multiple batches or experiments by extending the functionality of CytoNorm [30] batch alignment algorithm. Furthermore, we provide a means to transfer labels from reference datasets onto new datasets using machine learning classification algorithms, allowing for reproducible analytical workflows across multiple experiments. An extension of this process allows for the analysis of very large datasets, where the size of the dataset may exceed the memory capacity of the computer being used. Analysis with Spectre can be applied to flow, spectral, and mass cytometry datasets, consisting of tens to hundreds of millions of cells, as well as to scRNAseq data and HD imaging data (such as Imaging Mass Cytometry, IMC). Spectre is available as an R package or Docker container from Github (https://github.com/immunedynamics/spectre).

Here we demonstrate the utility of the package by analyzing cells isolated from murine bone marrow (BM), spleen, and central nervous system (CNS) following inoculation with West Nile virus (WNV), measured by HD flow, spectral, or mass cytometry (CyTOF). Because of the significant cellular dynamics that occur in multiple tissues in response to infection with WNV, this provides an ideal model for demonstrating analysis workflows using Spectre.

## METHODS

### Sample preparation and acquisition

Ethical approval for the experimental use of mice was obtained from the Animal Ethics Committee at the University of Sydney. Briefly, mice were anesthetized with 250-300 μL of Avertin anesthetic via an intraperitoneal injection, and then inoculated intranasally (*i.n*.) with 10 μL of PBS or a lethal dose of West Nile virus (WNV) in sterile PBS (6×10^4^ plaque-forming units [PFU]). After 1-7 days post infection (*p.i*.), mice received 350 μL of Avertin anesthetic, followed by vena cava section and left ventricular cardiac perfusion with 10 - 30 mL of ice-cold PBS. Spleen, BM and CNS tissue were then extracted and prepared for flow/spectral [33] or mass cytometry [32] as previously described. Labelled samples were acquired on a 5-laser BD LSR-II, a 10-laser custom BD LSR-II, a 5-laser Cytek Aurora, or a Fluidigm Helios. Initial cleanup gating and compensation was performed using FlowJo v10.7, where single live leukocytes were then exported as CSV (scale value) or FCS files.

### Data management in R via data.table

Data management and operations within Spectre were performed using the *data. table* format, and extension of R’s base *data.frame*, provided by the *data.table* package [34]. This table-like structure organizes cells (rows) against cellular features or metadata (columns). This simple *data.table* structure allows for the high-speed processing, manipulation (subsetting, filtering, etc.), and plotting of large datasets, as well as fast reading/writing of large CSV files. While CSV files are the preferred file format in this context, Spectre also supports reading and writing FCS files through the use of *flowCore* [35], where the data are then converted from an FCS file into a *flowFrame*, and then into a *data.table*.

### Data pre-processing and transformation

In Spectre, the entire dataset to be analyzed is condensed into a single *data.table*. Each row (cell) contains cellular expression data and metadata pertaining to the file, sample, group, and batch, stored in separate columns. Metadata for each of the columns (e.g., alternative cellular marker names, etc.,) in the *data.table* can be stored separately, if required.

In order to reduce the contribution of background to measured signal, and to convert cytometry data into a linear space, we facilitated ArcSinh transformation of data using the *do.asinh* function in Spectre. This function allows a transformation co-factor to be specified for all columns (typical in mass cytometry) or for individual columns (typical in flow or spectral cytometry). The resulting transformed values are added to the dataset as new columns. This addition of new values, rather than replacement of original values, enables greater data retention and transparency. Additional pre-processing steps, such as noise reduction for values below zero, and normalization of values onto the range of zero to one, are provided in the *do.noise.reduce* and *do.normalise* functions respectively, which similarly adds columns of new data.

### Batch alignment and integration

To facilitate batch alignment in Spectre, we developed a wrapper for CytoNorm [30] using the *do.align* function. CytoNorm utilizes reference samples (aliquots of a control sample run with each batch) to determine batch-derived technical differences and calculates a quantile conversion model to align data from each of these batches. This conversion model is then applied to samples in each batch, removing technical variation while preserving biologically relevant differences. The *do.align* function operates on a *data.table*, and allows for reference and target data to be easily specified in a dataset containing a mixture of batches. Quantile conversions for the entire dataset, or conversions for individual clusters, can then be calculated, resulting in a new set of ‘aligned’ features that are compatible across the batches. Since raw, ArcSinh transformed, and aligned data are preserved in the *data.table*, transparency of these analytical processes is maintained throughout the analytical process.

### Clustering of cells/events

Clustering was implemented using the *run.flowsom* function, which acts as a wrapper around the FlowSOM function (available as ‘FlowSOM’ from Bioconductor [36]), which organizes acquired events (i.e., cells) into clusters using a self-organizing map (SOM), and then groups these clusters together into ‘metaclusters’. The function accepts a *data. table* and set of column names for generating clusters as input. Once clustering is performed, *run.flowsom* returns the *data. table* with new columns containing the cluster and metacluster IDs for each cell. *Spectre’s run.flowsom* provides two options for specifying a target number of metaclusters. By default, FlowSOM will determine the number of metaclusters to generate by performing consensus hierarchical clustering on the clusters using a range of different metacluster numbers. It then computes the variance in each metacluster number and determines the point in which the variances suddenly decrease at a gentler rate (the elbow point), which is regarded as the optimal number of metaclusters for the dataset. We refer the reader to the FlowSOM publication for more details on how the optimum number is chosen [5]. Alternatively, users can manually specify a target number of metaclusters (which is recommended over automatic determination) or chose not to generate metaclusters at all. In addition to the ability to choose the number of metaclusters to create, the user is also able to change the SOM grid size (*xdim* and *ydim* parameters), the seed used to generate clusters and metaclusters for reproducibility.

### Dimensionality reduction

Non-linear dimensionality reduction (DR) was implemented in Spectre using the *run.tsne*, or *run.umap* functions, which are wrappers around t-SNE (available as ‘rtsne’ from CRAN [37]), and UMAP (available as umap from CRAN [38]), respectively. Both the *run.tsne* and *run.umap* functions accept a *data.table* and set of column names for DR as input, and return the *data.table* with new columns containing tSNE or UMAP coordinates for each cell. We also provide a function for running Principal Component Analysis (PCA), a linear DR approach, using the *run.pca* function, acting as a wrapper around *prcomp* (available in the ‘stats’ package from CRAN [39]). The input data for *run.pca* can either be individual cells (to find cell markers that contribute to the variance), or individual samples/patients with summary data (such as median fluorescence intensity (MFI), cell counts or proportions). The output of *run.pca* is similar to *run.tsne* and *run.umap*, though additional outputs can be generated, including scree and contribution plots for detailed assessment of the source of variance in the dataset.

### Plotting and visualization

Plotting of cellular data was implemented in the *make. colour.plot* function, serving as a wrapper around ggplot2 [40]. The dataset, and desired columns to use as X and Y axis are taken as input. By default, each plotted cell is colored by relative plot density. Where desired, each cell can be colored by the level of expression of a specified marker (column), or color by some factor (e.g. cluster, sample, or group etc). To reduce the effect of outlier datapoints on the color scale, minimum and maximum thresholds are applied by default as 0.01 (1^st^ percentile) and 0.995 (99.5^th^ percentile) respectively, so that datapoints with values below or above these thresholds are colored at the minimum or maximum respectively. These can be modified where required. In cases where a subset of the dataset is being plotted (e.g. separate samples or groups, etc) the limits of the X, Y, and color parameters can be set by the complete dataset, allowing for the plots to be compared directly. By default, plots will automatically be saved to the current working directory, allowing for the rapid generation of plots. This plotting functionality is extended in the *make. multi.plot* function, which arranges multiple plots consisting of set of colored plots (e.g. for examining marker expression) or set of sample group divisions (e.g. for comparing changes between groups).

### Aggregation and summary data

A number of analysis steps require the aggregation or summary of data, either by sample, by cluster, or by both. To facilitate this, we implemented two functions – *do.aggregate* and *write.sumtables*. The *do.aggregate* function takes advantage of the fast aggregation function *of data.table*, providing the mean or median level of expression of each marker per cluster/population. This is helpful for plotting expression heatmaps, allowing for the interpretation of cluster/population identities. For every population in each sample, the *write.sumtables* function will calculate the percentage of each population as a percentage of cells in each sample (as well as cell counts for each population per sample if a total count of cells per sample is provided). Additionally, for all specified cellular markers, the median expression level, as well as the percentage of each population that expresses the marker are calculated for each population in every sample. These summarized *data.tables* are saved to disk as CSV files, in the format of samples (rows) vs features (columns), which can then be used for quantitative and statistical analysis, either in the same workflow, or in a separate pipeline.

### Heatmaps

To facilitate the generation of heatmaps for examining marker expression on populations, or for comparing populations across samples, we implemented *‘make.pheatmap’* as a wrapper around the pheatmap function from pheatmap [41]. In *make.pheatmap*, the user must specify the dataset, the column that contains values to be used as the heatmap rows (e.g. sample or cluster identity), and the columns to be plotted as heatmap columns. This can be used to examine expression of marker (columns) on each cluster (rows), or can be used on z-score transformed data plotting immune features (columns) vs samples (rows), to reveal data variance for each cellular feature being measured.

### Group and volcano plots

To facilitate quantitative and statistical analysis, we implemented functions to create graphs (*make.autograph*) and volcano plots (*make.volcano.plot*). Graphs are constructed using the *ggplot2* R package, and integrate pairwise statistical comparisons between groups using functionality from the *ggpubr* R package [42]. Pairwise comparisons between pairs of groups are performed using either Wilcox test (equivalent to a Mann-Whitney test), or a t-test. Overall group variance is assessed using a Kruskal Wallace test, or a one-way ANOVA. Volcano plots were generated using the *EnchancedVolcano* package from Bioconductor [43], and customized to work seamlessly with *data. table* data structures. Initially the relative −fold change of values in each immune parameter between two groups are calculated, along with the p-value of that change. Each immune parameter can then be plotted as fold change vs p-value, allowing for a rapid assessment of changes with strong and significant effects between groups.

### Classification

Following batch alignment, the transfer of population labels from one dataset to another was implemented via two functions: *run.train.knn.classifier* and *run.knn.classifier*. The *run.train.knn.classifier* function uses the *caret* [44] package to train a k-nearest neighbor (kNN) classifier [42] using the labelled dataset, allowing for the identification of the two most closely-matched cells across each dataset. It first normalizes each feature (marker) of the dataset to a range between 0 and 1 using the *preprocess* function of *caret*. This is to ensure a uniform distribution of values across all features. Thereafter it trains the KNN classifier using the n-fold cross-validation (n-fold CV) technique. The n-fold CV technique splits the data into n complementary subsets and uses n-1 of those to train the classifier (training data), and the remaining one to test the effectiveness of the trained classifier (testing data). This process is repeated n times, whereby in each iteration different subsets of data are used for training and testing data. For n-fold CV, the overall quality of the classifier is computed as the average quality obtained for all n rounds of training and testing. The *run. knn. classifier* function then runs a kNN classifier provided by FNN R package [45] on a *data. table* without cellular population labels, returning the *data.table* with a new column of population labels added.

## RESULTS

### Spectre facilitates comprehensive end-to-end integration and analysis of large high-dimensional cytometry datasets

Spectre was developed to facilitate rapid and flexible analysis of large and complex cytometry datasets across multiple batches or experiments. Specifically, Spectre facilitates data pre-processing (Figure 1A), alignment of data from multiple batches/experiments (Figure 1B), clustering (Figure 1C), dimensionality reduction and visualization (Figure 1C), manual or automated population classification and labelling (Figure 1C), as well as extensive plotting and graphing options for qualitative and quantitative statistical analyses (Figure 1D). Key to this process is strategic selection, implementation, and customization of high-performance computational tools; and the development of wrapper functions around these tools, enabling them to operate on and produce the same data format for input and output, respectively. This allows for multiple analysis and plotting/graphing tools to be seamlessly weaved together into single analysis workflows, where functions can be used throughout the stages of the workflow, drastically increasing ease of use. Moreover, this modularity and flexibility allows these workflows to be adapted to meet different experimental requirements or analytical approaches. Spectre can be used on data generated by a variety of single-cell technologies, including flow, spectral, and mass cytometry (Figure 1E). In addition, following some additional pre-processing, Spectre can also be used for the analysis of data generated by scRNAseq or HD imaging technologies such as IMC. (Figure 1E).

**Figure 1.**
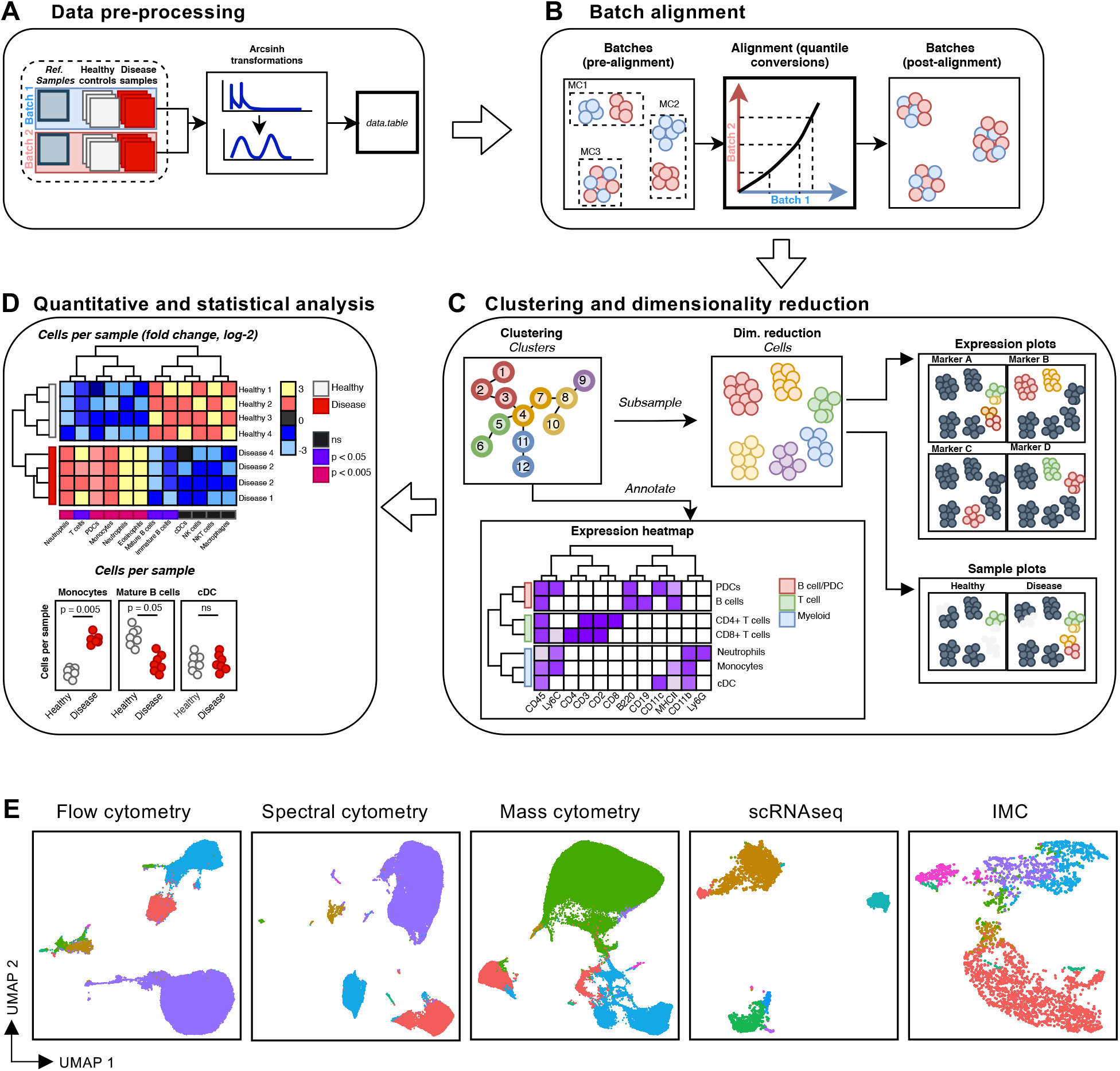
Spectre analysis overview. An overview of Spectre’s analysis workflow. A) Data preparation steps including sample, group, and batch annotation, in addition to ArcSinh transformation. B) Batch alignment using CytoNorm. C) Clustering and dimensionality reduction, along with marker expression plotting and expression heatmaps. D) Quantification and statistical analysis through z-score/fold-change heatmaps and grouped dot plots. E) Application of Spectre’s analysis workflow to data generated by different technologies, including analyzing a split murine spleen sample by flow cytometry, spectral cytometry, and mass cytometry. Also shown are unrelated PBMC data analyzed by single-cell RNA sequencing (scRNAseq) acquired from 10X genomics, via the Seurat webpage (https://satijalab.org/seurat/v3.2/pbmc3k_tutorial.html) and imaging data generated by Imaging Mass Cytometry (IMC).

### Data management and pre-processing

Critical to our approach is the use of the *data. table* data structure and operations, in place of prevalent FCS/flowFrame objects or other data structures. The *data.table* format is an enhancement of base R’s *data.frame*, a table-like structure commonly used to store data. This enhancement allows for fast and efficient aggregation and manipulation of data through the use of concise flexible syntax and low-level parallelism [34] (Supp. Figure 1). Using Spectre, all samples in the analysis are merged into a single *data.table*, with relevant sample, group, and batch information stored in separate columns. As a result, each row contains all the information relevant for a particular cell, making data manipulation and filtering with *data. table* simple and fast.

A key initial step in computational analysis is the transformation of cellular expression data. Biologically meaningful results are most easily interpreted through plotting cellular expression on a logarithmic scale. Because of potentially misleading visual artifacts for signals at the low end of the logarithmic scale, the logicle/bi-exponential scale was developed, where the high end of the scale is logarithmic and the low end of the scale is converted into a linear scale, and the scale then returns to logarithmic at values below the linear component (Supplementary Figure 2A) [46, 47]. Critically, this allows for the compression of low-end data points with high spreading error, autofluorescence, or noise into a linear space around zero, which can be tailored for the requirements of each channel (Supplementary Figure 2B). For computational analysis to meaningfully manage biological data, a similar compression of low-end data needs to be performed. In cytometry, this is commonly performed using the ArcSinh (http://mathworld.wolfram.com/InverseHyperbolicSine.html) transformation [48]. The data values are transformed into a linearized format, where compression of low-end values is determined using a specified co-factor to determine the extent of compression around zero (Supplementary Figure 2B).

Using Spectre, ArcSinh transformation was applied to data from each channel/marker individually, using different co-factors, allowing for highly customizable data transformations. For flow, spectral, and mass cytometry data, we tested various co-factor values across multiple channels (Supplementary Figure 3-4). Overall, we found that a co-factor of 5 to 15, was suitable for all mass cytometry channels. However, in our experience, we found a range between 100 to 10,000 to be suitable for different channels in conventional or spectral flow cytometry data.

Because low-end data compression through ArcSinh transformations may still result in significant spread of the negative population around zero, we reasoned that this could be reduced by converting any values that occur below zero, to zero, effectively reducing the contribution of noise below zero. This was implemented in the *do.noise.reduce* function, where a minimum threshold can be set, where all values below that value will be converted to that value (Supplementary Figure 5). Additionally, we developed a function for re-distributing (aka ‘normalizing’) ArcSinh data between two new values, usually 0 and 1. This prevents markers with extremely high expression levels from exerting greater influence over clustering and DR results, when compared to other markers. This was implemented in the *do.normalise* function (Supplementary Figure 5). Although we have demonstrated the utility of the *do.noise.reduce* and *do.normalise* functions in Supplementary Figure 5, they were not used for the rest of the data presented in this paper.

### Integrating data from multiple batches or experiments into a single feature space

When samples are prepared, stained or run in multiple batches, technical batch-effects can occur, usually consisting of shifts in signal intensity in one or more markers. Because of this, data clustered together with uncorrected batch effects may separate samples based on the batch they belong to, a confounding factor that substantially hinders aggregated analysis of datasets from multiple batches or experiments. To provide a comprehensive and adaptable batch alignment and data integration approach, we expanded on the functionality of CytoNorm within Spectre. Users specify reference control samples that Spectre uses to determine the alignment conversions. Typically, these are aliquots of a single ‘healthy’ patient sample that are run with each batch of samples. These reference samples should span the full range of the data seeking to be aligned. Where marker expression data is absent on the reference controls (e.g. absence of activation markers on cells from healthy donors), alignment is not feasible, but the original expression data for those markers can still be analyzed. In cases where multiple aliquots of a single control sample are not available, multiple control samples of the same type (e.g., healthy mouse BM) that are run with each batch can be used as the next best. Finally, it is also possible to use all samples from each batch as ‘reference’ controls, taking the entire range of marker expression from each batch into account. However, these later approaches should be used with caution, as any biological differences between the samples used for reference will be interpreted as technical variation due to batch effects. To execute this alignment process in Spectre, user-indicated reference samples are extracted from the combined *data.table* (Figure 2A) and clustered using FlowSOM. For each resulting metacluster, quantile conversion coefficients between each batch for each marker are calculated (Figure 2B). Cells from the full dataset are then mapped to the FlowSOM grid, and each cell is assigned to its nearest metacluster (Figure 2C). Quantile conversion is then applied to the cells in each metacluster, unifying cells from each batch into a single feature space, while preserving biological differences between experimental groups (Figure 2C).

**Figure 2.**
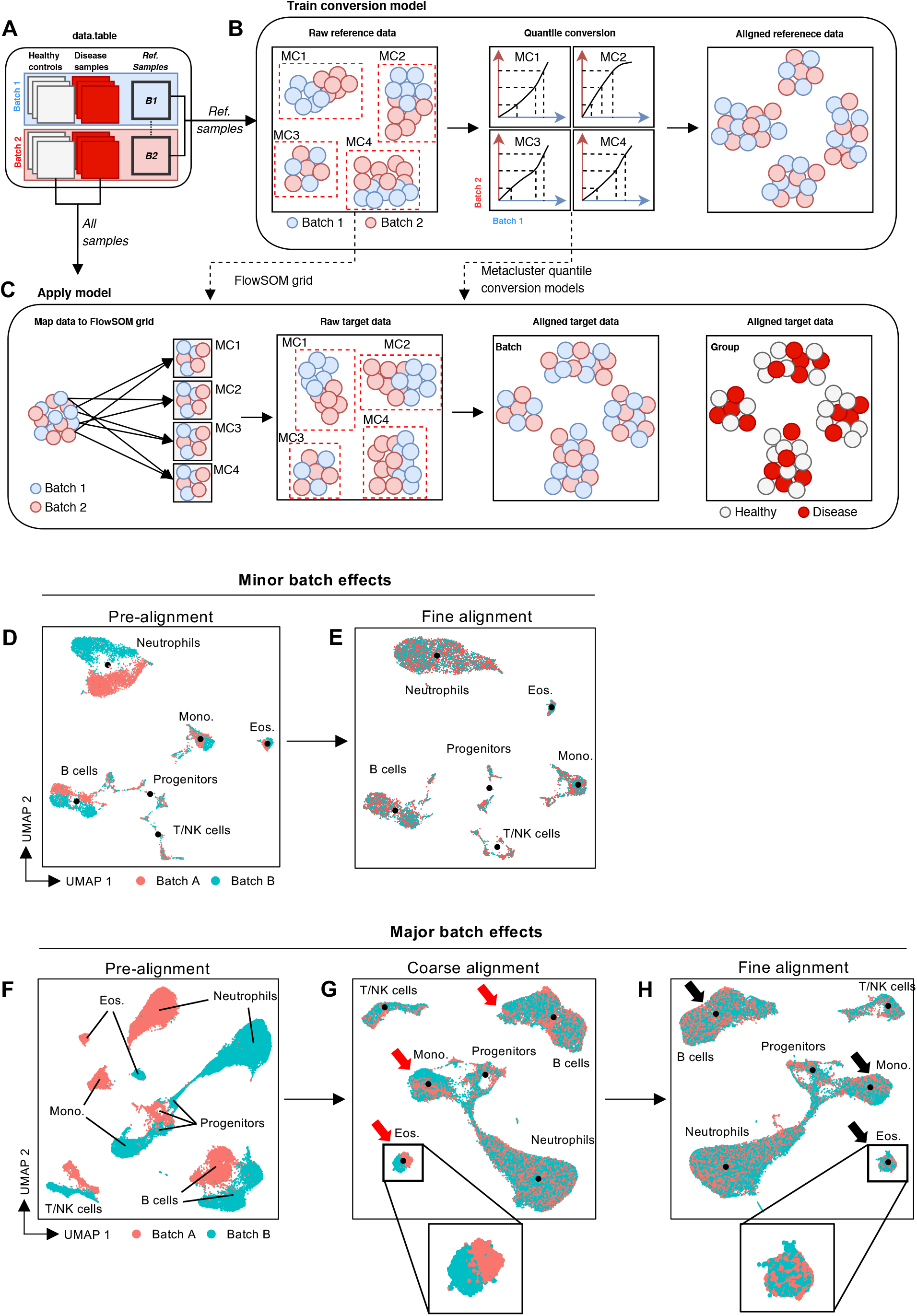
Batch alignment using CytoNorm. Batch alignment process using CytoNorm. A) Reference samples acquired with each batch are extracted from the *data. table*, and B) clustered using FlowSOM. Metacluster-specific quantile conversion models are then calculated. C) Cells from all samples/batches are mapped to the FlowSOM grid, and assigned to their nearest metacluster. Cells are then aligned using the metacluster-specific quantile conversion models D) Two sets of BM samples with synthetic batch effects introduced in a population-specific manner. E) CytoNorm alignment was performed using a metacluster-specific alignment process (fine alignment). F) Two sets of BM samples generated with slightly different panels, but targeting the same cellular markers, resulting in significant batch effects. G) CytoNorm is initially performed on the whole dataset (coarse alignment) by mapping the entire dataset into a single metacluster, where H) subsequent FlowSOM clustering allowed for further metacluster-specific alignment (fine alignment).

To verify the robustness of this process, we applied batch alignment to a set of mock- or WNV-infected BM samples, where synthetic data manipulations were introduced in a population-specific manner to half the samples, to mimic population-specific batch effects. One healthy BM sample from each ‘batch’ were selected as a reference controls, aggregated together, and plotted using UMAP, revealing subsets of neutrophils, eosinophils, monocytes, B cells, T/NK cells and progenitors. Batch-specific differences in their distributions were evident in the UMAP plots (Figure 2D). When FlowSOM was run, consistent populations from each batch were captured within the same metacluster, despite the presence of batch effects. As such, metacluster-specific alignment with CytoNorm was able to adequately integrate the cells from each population into a unified dataset (Figure 2E).

To test this process on datasets with more substantial batch-effects, we applied batch alignment to two sets of mock- or WNV-infected BM samples, prepared and acquired months apart, using slightly different panels. After matching column names between the two datasets, the disparity of these two datasets was apparent upon plotting with UMAP (Figure 2F). In this scenario, the batch effects were so significant that cells from the same population in each batch did not map to the same metacluster, thus preventing alignment. To address this, we initially performed a ‘coarse’ alignment by mapping all cells from each batch into a single metacluster, and performing quantile alignment on the entire dataset. This corrected the majority of batch effects, though some metacluster-specific effects were still evident (Figure 2G, inset). Nevertheless, the resulting data was sufficiently aligned so that populations from each batch could now be mapped into consistent metaclusters (Figure 2G), allowing for a more fine-tuned alignment (Figure 2H) that corrected residual metacluster-specific batch effects (Figure 2H, inset) and unified the dataset into a single feature space.

### Clustering and dimensionality reduction strategies

A critical analytical step in HD cytometry is the comprehensive identification of cellular populations, including those that are well-established and those that are yet to be characterized. This is particularly relevant when comparing between experimental groups (e.g., diseased patients compared to healthy controls). Clustering and dimensionality reduction are powerful techniques for identifying cell populations and contrasting them between groups. Clustering tools collect phenotypically similar cells into groups (clusters) in a data-driven fashion (i.e., cells are grouped together based on the similarity of their marker expression). The output of many clustering approaches are plotted as a collection of nodes, where each node represents a cluster that contains a number of cells. These nodes are typically connected by a form of minimum spanning tree (MST) [5] and colored by mean or median marker expression of the cells within each cluster. However, verifying the phenotypic heterogeneity (or homogeneity) of cells captured within a cluster is difficult when looking at the data at the cluster level. An alternative approach is to compress cellular data onto two dimensions using non-linear dimensionality reduction tools such as t-SNE [7, 8] and UMAP [9], and to visualize these on a scatter plot, coloring each cell by marker expression level or cluster/metacluster ID.

Spectre supports clustering and DR by providing a wrapper function for FlowSOM, t-SNE, and UMAP. These functions accept and return data in the *data.table* format. Whilst clustering tools such as FlowSOM scale well to large datasets [49], DR approaches such as t-SNE and UMAP do not; as they incur lengthy computing time, excessive memory usage, and significant crowding effects that inhibit their utility (Supplementary Figure 6). Whilst some improvements to runtime (flt-SNE [50]) and plot crowding (opt-SNE [51]) have been made, scalability and plot crowding limitations persist. As DR tools are primarily used to visualize cellular data and clustering results, we propose plotting only a subset of the clustered data, which addresses scalability and retains legibility. By using proportional subsampling from each sample, the relative number of cells from each cluster in each sample can be preserved in a smaller dataset, allowing for interpretable analysis via DR. Putative cellular populations amongst the clusters can then be identified, and annotated in both the subsampled DR dataset, as well as the whole clustered dataset. The whole annotated dataset can subsequently be used in downstream quantification and statistical analysis.

To demonstrate this strategy, we applied clustering and DR to a dataset of brain cells from mock- or WNV-infected mice (Figure 3A). The whole dataset was clustered using FlowSOM (Figure 3B), and the clustered data were proportionally subsampled for plotting with UMAP (Figure 3C). In this case, the number of cells extracted from each sample were in proportion to the total cells recovered from each brain, ensuring that DR plots accurately reflected sample composition (Figure 3D). By examining expression heatmaps (Figure 3E) and colored DR plots (Figure 3F) we manually determined cluster population identities, and these annotations were applied to both the subsampled and full datasets.

**Figure 3.**
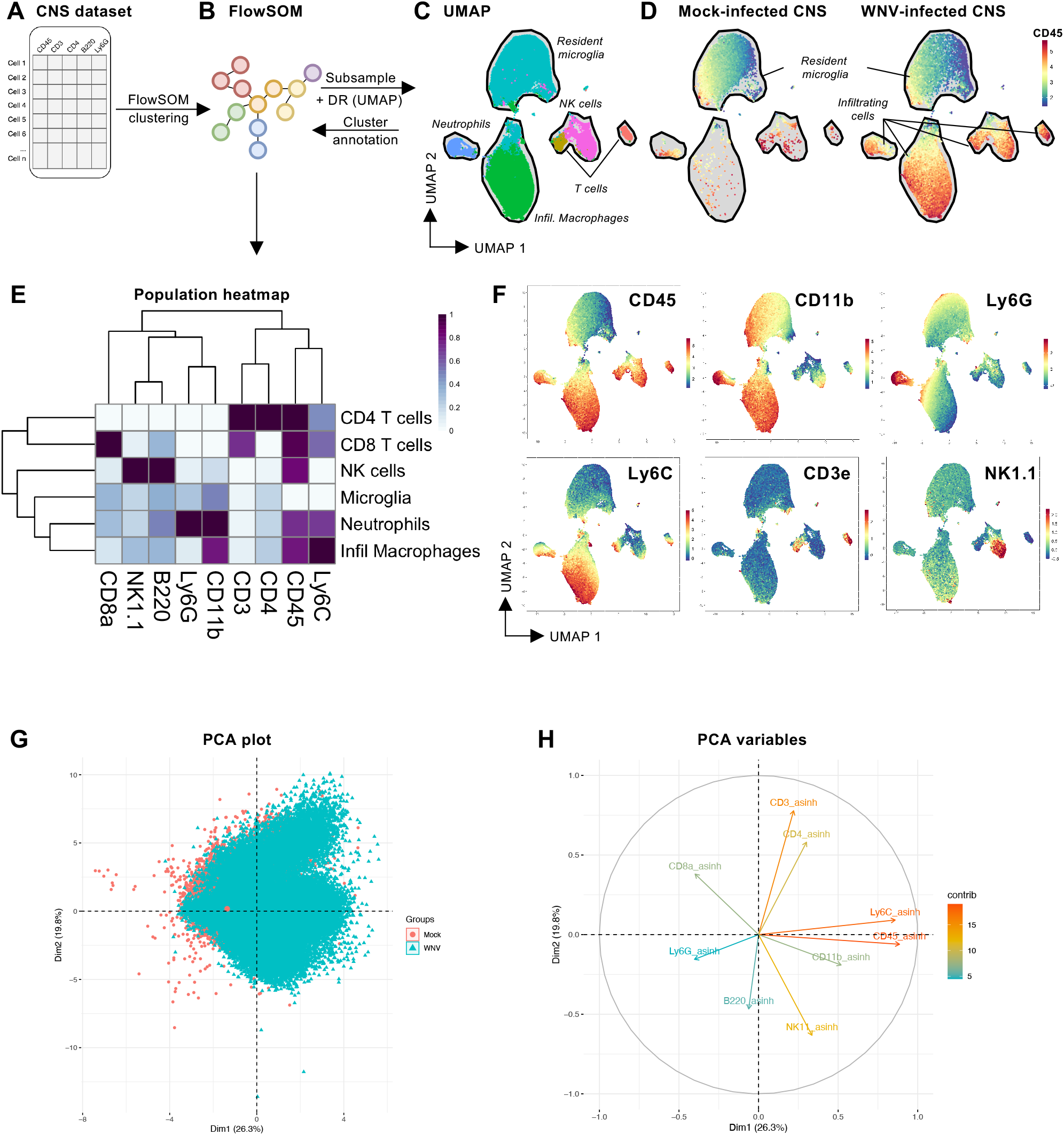
Clustering and dimensionality reduction using Spectre. A) A dataset of cells isolated from mock- or WNV-infected CNS were used to demonstrate clustering and DR in Spectre. B) FlowSOM clustering was performed on the full dataset, which was C) then subsampled and plotted using UMAP. D) Parsing the dataset by each experimental group reveals substantial changes to immune populations. E-F) An examination of marker expression on each cluster allows for a user-determined annotation into biological relevant cell types. G) Analysis using PCA allows for visualization of the data variance, and H) the relative contribution of markers to the first two principle components.

The choice of markers used to inform the generation of clusters/DR results is dependent on the overall goal of analysis. Cellular markers may be broadly categorized into two groups: static (stably expressed) or dynamic (changing expression). Typically, statically expressed markers are helpful for identifying consistent cell *types* (e.g. T cells, B cells, etc.), whereas dynamic markers are helpful for identifying reactive cellular *states* (e.g. activated, resting, etc.). When seeking to discover novel cellular populations or states, incorporating all cellular markers may be of benefit, as both stable cell types and dynamic cellular states will be captured in separate clusters. However, a more selective approach may also be desired, such as using statically expressed markers to capture known populations within clusters, and then examining those stable populations for dynamic changes in activation status. Along with domain-specific knowledge about the expression patterns of various markers, it is possible to identify markers that contribute most to the level of variance across a given dataset. To do this we can using principal component analysis (PCA), (Figure 3G-H) to determine the relative contribution of each cellular marker to the overall variance of the dataset.

### Multi-level immune profiling

Many populations of interest, such as hematopoietic stem cells (HSC) in the BM, are of very low frequency within individual samples. As such, representation of these rare subsets may be extremely sparse on DR plots, relative to more abundant populations. Moreover, more nuanced clustering and analyses of such populations are often desired, but the global data structures provided by more abundant subsets of cells may dominate the analysis. To address this, we extended our analysis approach to enable the exploration of data at multiple levels. Upon clustering the complete dataset, clusters representing rare or novel populations can be isolated and re-clustered independent of other populations and thereafter annotated in greater detail. Expanding on this approach, multiple lineages can be split and profiled independently, then re-merged, retaining detailed cluster annotations for combined plotting and quantitative analysis. This process is conceptually similar to hierarchical approaches such as hierarchical-SNE (h-SNE, [52]), but accommodates a more bespoke tailoring of the analytical process.

To demonstrate this, we examined a dataset of BM HSCs generated by mass cytometry. Clusters containing HSC and progenitors (denoted by CD117 expression) (Figure 4A) were extracted from the full dataset, and subjected to independent clustering and DR (Figure 4B). This independent analysis allowed for a more detailed assessment of low frequency subsets (Figure 4C) that were not easily assessed in the plots from the full dataset.

**Figure 4.**
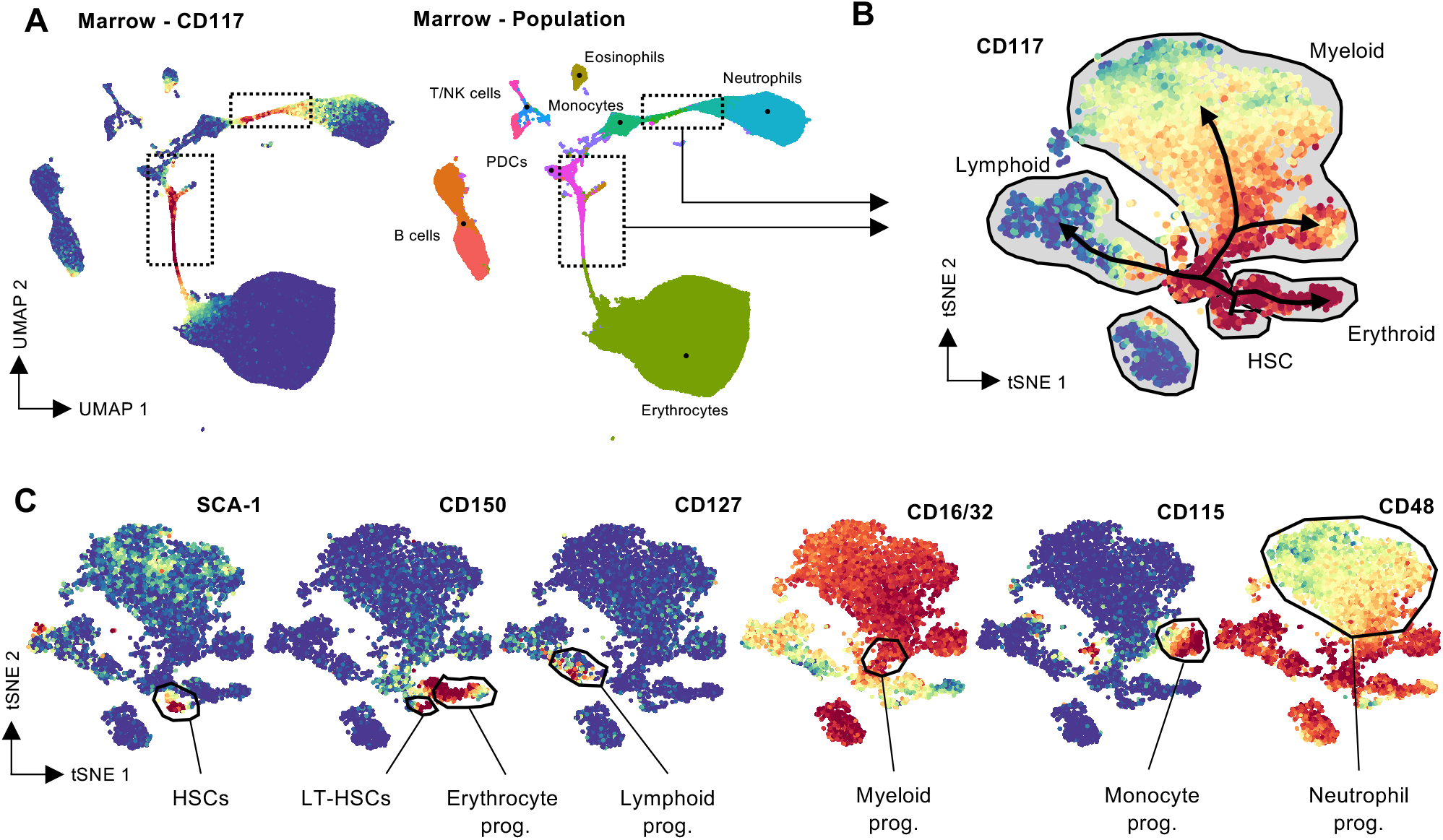
Multi-level analysis for profiling of rare populations. A) A UMAP plot (left) where clusters representing stem cell and progenitor subsets were identified via CD117 expression. Through cross-referencing against FlowSOM clusters (right), these cells were B) subjected to new clustering, subsampling, and plotting using UMAP. C) Expression color plots reveal low frequency cellular subsets that were difficult to otherwise detect on the full UMAP plot.

### Automated cellular classification and label transfer between aligned datasets

A crucial application of computational analysis is *discovery* - defining novel subsets and/or investigating experimental changes in novel subsets or states in new diseases, tissues, or experimental conditions. Analytical approaches to this often depend heavily on unsupervised techniques, such as clustering and DR. Such analyses culminate in an annotated datasets in which each cell is manually assigned a population label by the user. Alternatively, other studies seek to apply a more *repetitive* analytical process, using semisupervised tools, such as automatic gating, to replicate a method of population identification over a large number of samples. While effective, many of these tools rely on some form of gating strategy and forgo the use of any unsupervised techniques to expedite gating, thereby limiting the possible identification of new or complex/overlapping populations. In contrast, machine learning-based approaches provide the opportunity for automated transfer of cellular labels from an annotated dataset to a novel dataset, following alignment of the datasets into a single feature space (as demonstrated in Figure 2). To facilitate this, we provided functions within Spectre to train and run classifiers, a type of machine learning approach designed to predict the class of given data points. As opposed to clustering, which groups cells together based on marker expression, classifiers assign cells a label based on ‘training’ data. This training data could be previously gated, clustered, or annotated and is used by the classifier to determine how given features (marker expression) relates to a class (cluster or cell phenotype).

Inspired in part by Seurat’s mutual nearest neighbors approach for data integration [19, 20], we reasoned that a simple nearest neighbors approach could facilitate rapid label transfer between datasets. In Spectre, we implemented a k-Nearest Neighbor (kNN) classifier, which classifies unlabeled cells based on the label of their K^th^ nearest neighbor within the training dataset. To determine the accuracy of the kNN classifier, we divided a labelled dataset into two halves (Supplementary Figure 7A) - one half retained cellular labels and served as the ‘training’ dataset, and the other half had their cellular labels hidden, and served as the validation dataset. The kNN model was trained on the labelled data and applied to the unlabeled data (Supplementary Figure 7B), to compare against their original cellular labels (Supplementary Figure 7C). Generally, a kNN classification scheme with *k* = 1 tended to predict cellular labels between datasets with high accuracy (> 98.6%), although we found that increasing *k* here marginally increased this accuracy to over 98.9% in this case (Supplementary Figure 7D).

To demonstrate the kNN classification process in an experimental context, we aligned two datasets of BM cells using CytoNorm (shown in Figure 2D-E), where the first dataset (batch A) contained labelled cells, and the second dataset (batch B) did not (Figure 5A). Following alignment, the data was split into the labeled and unlabeled datasets (Figure 5B), where the kNN classifier was trained on the labelled cells (batch A) and applied to unlabeled cells (batch B) with *k* = 1 (Figure 5C). When we compared the predicted cellular labels to manually annotated cellular labels, we found the classifier was able to accurately transfer cellular labels between the two datasets (Figure 5D). Merging of the two datasets the resulted in a fully annotated dataset (Figure 5E).

**Figure 5.**
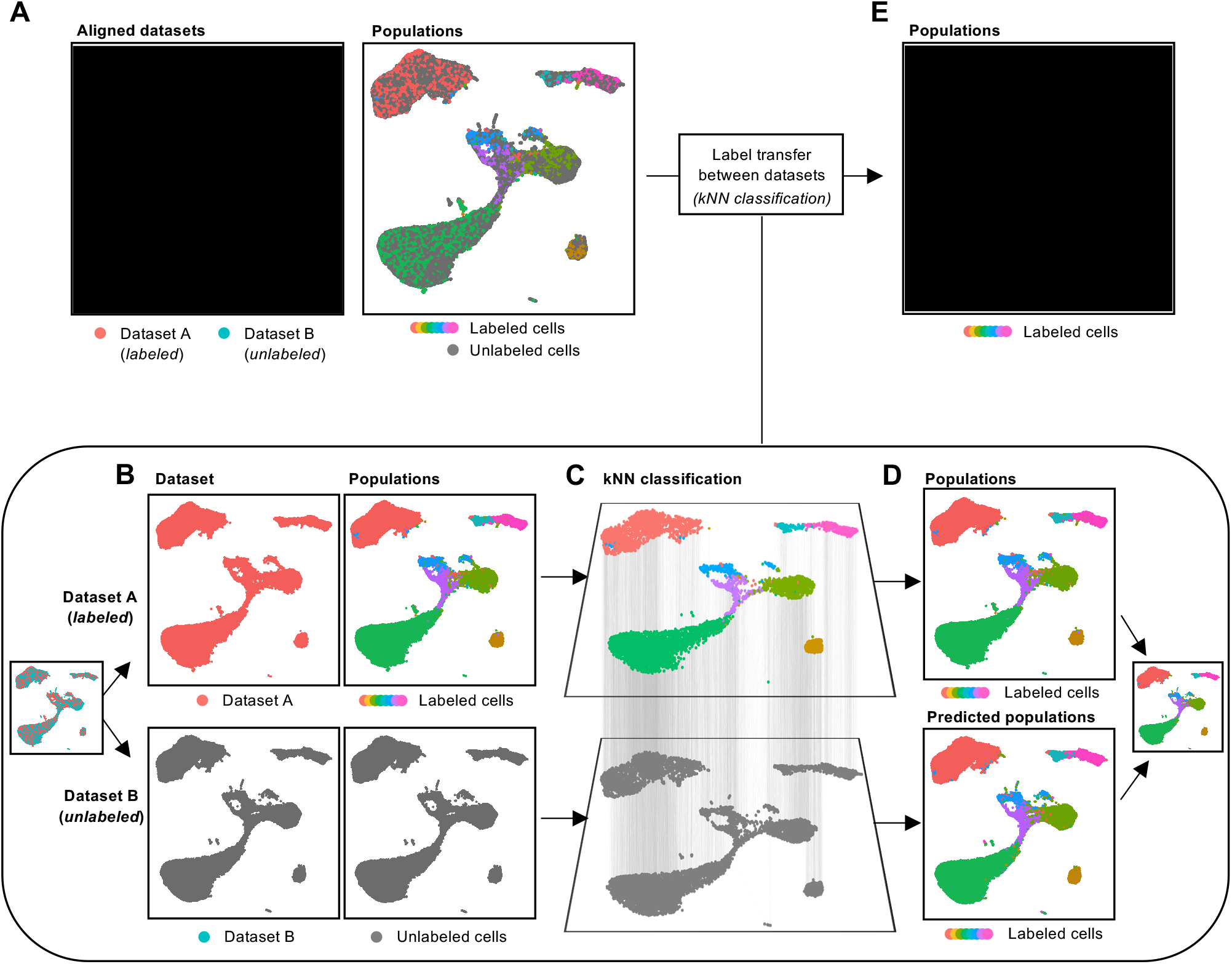
Cellular classification and label transfer. A) A dataset of two batches of BM cells, following alignment with CytoNorm. One batch contains annotated clusters, and the other does not. B) The dataset is split into each batch (labeled and unlabeled), and a kNN classifier trained on the labelled dataset with k = 1. C) The kNN classifier was then applied to the unlabeled dataset, D) resulting in an accurate transfer of cellular labels between datasets.

### Quantitative and statistical analysis

The endpoint of most analytical workflows is to make quantitative and statistical comparisons between experimental groups. Many components in this workflow can be automated to simplify the processing steps and reduce the time taken to generate relevant statistics and plots. To facilitate this, we have developed a series of functions to summarize a dataset rapidly at either the cluster or cellular population level, resulting in a series of summary tables. For each population in each sample, these tables summarize the proportion of cells, total cell counts, marker expression levels, and proportion of cells that are ‘positive’ for each cellular marker, which can then be used to generate quantitative plots. The generation of grouped scatter or violin plots (Figure 6A) provide a simple method to assess changes of a single feature (e.g. number of infiltrating macrophages per brain) between experimental groups, including grouped or pairwise statistical comparisons. However, more global statistical analyses are often desired. The generation of z-score heatmaps (Figure 6B) provide an overview of relative changes between samples, with optional clustering on samples (rows) or features (columns) based on similarity. Additionally, the result of pairwise comparisons between groups can be indicated for each heatmap column, revealing statistical significance for uncorrected or false discovery rate (FDR)-corrected p-values. Furthermore, PCA plots (Figure 6C) and volcano plots (Figure 6D) provide a further global view of how these immune features differentiate samples within the experimental context.

**Figure 6.**
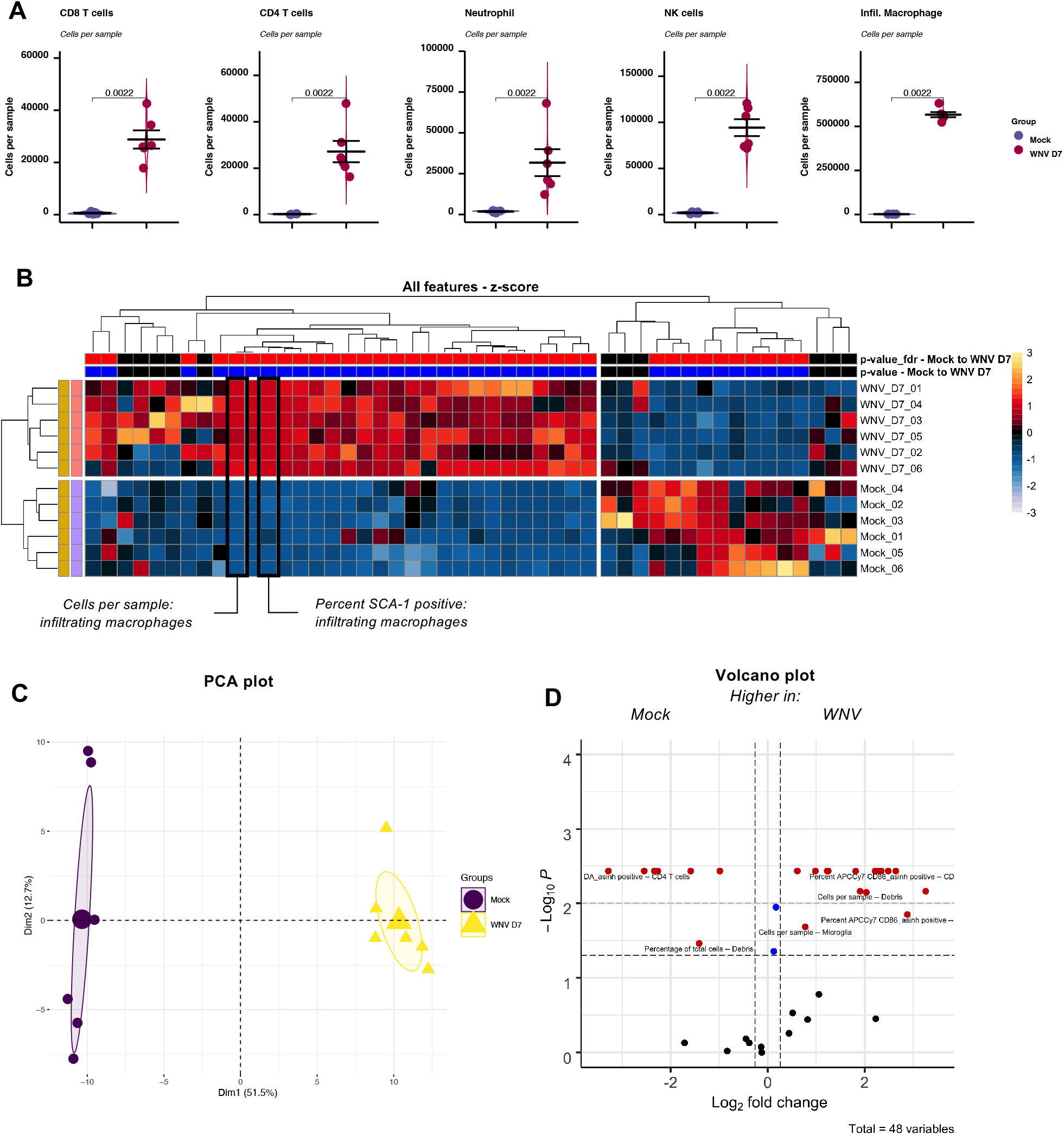
Quantification and statistical analysis. Quantitative and statistical analysis of CNS samples. A) Grouped dot plots with underlying violin plots indicating the relative cells per sample for each population in the dataset. Pairwise statistical analysis for non-parametric data was performed using a Mann-Whiteny/Wilcox test, where p < 0.05 was considered significant. Importantly, data for each plot were not adjusted for multiple comparisons. B) The z-score for each column of data was calculated, and plotted using the make.pheatmap function. Each row represents a sample, and each column represents a measured immune feature. Rows and columns were clustered using euclidean distance. Pairwise comparisons between experimental groups were calculated using a Mann Whiteny/Wilcox test, and the significance results plotted as significant (p < 0.05) or non-significant (p > 0.05) for each column. To adjust for multiple comparisons, p-value results were corrected using a FDR correction, and results plotted as significant (p < 0.05) or non-significant (p > 0.05) for each column. C) PCA results show the distribution of each sample, and D) volcano plots show the relative fold-change increase or decrease of each immune feature (X axis), and the inverse p-value of each change (Y axis).

## DISCUSSION

### Challenges in cytometric analysis

Historically, analysis of cytometry data involves a manual process of identifying cells, usually through drawing a gate on cells plotted with one marker vs another. As the number of features that can be measured on cells has continued to increase, the process of gating alone has become untenable for comprehensive analysis of large and complex datasets. This is in part due to the sheer number of gates that need to be drawn to fully parse a dataset, and in part due to the bias that is introduced by selectively including/excluding cells using one or two markers at a time sequentially. As a result, a number of computationally-driven approaches to data analysis have been developed, including automated gating, clustering, DR, and classification. These enable automated, data-driven data processing that may be performed in an unsupervised or semi-supervised fashion. However, these approaches come with their own limitations, including operational constraints, such as the number of cells that can be analyzed, given the available computational resources, the possibility of inaccurate identification of specific biologically-relevant populations in the dataset, and a lack of reproducibility between analytical runs (particularly when applying to new datasets). Critically, many analysis tools are developed as standalone packages that operate on specific data formats. This limits the interoperability of tools, especially when attempting to combine tools developed in different fields (such as cytometry and scRNAseq). As a solution, we developed Spectre: an adaptable and easy-to-use package for analyzing high-dimensional cytometry data. Spectre enhances existing computational tools through strategic implementation and customization of high-performance computational tools, and provision of wrapper functions to simplify and improve their flexibility and interoperability. This package expanded on many of the key aspects of the CAPX workflow [32] that has been utilized in a number of studies [53–57].

### As the foundation for data in Spectre, data.table allows for easy handling of large datasets

Many cytometry or single-cell analysis tools operate on custom data formats, such as the flowFrame [35], SingleCellExperiment [21], or Seurat objects [19]. Each custom data format may contain numerous elements, made up of both base R and custom data formats. Typically, primary cellular expression data is stored in a table or matrix, and separate elements contain metadata for each cell (cell number, sample, group, etc.,) or feature (marker names, parameters names, voltages, gene numbers, etc). Importantly, manipulated data (such as ArcSinh transformations or normalization) may be contained within the primary data table/matrix, or structured separately. The complexity of these custom formats, and the significant differences in structure between them, results in difficulties in converting one data format to another. While custom data formats may be convertible into other formats, including conversion to one of R’s base data formats, these conversions are non-trivial, and often result in the loss of important metadata. Of note, cytometry data is typically structured with rows representing cells, and columns representing cellular features, though this is transposed in scRNAseq data.

The foundation of Spectre is built upon the *data. table* package, which enhances R’s base *data.frame* format. The efficiency of *data. table* in performing basic data manipulation operations, such as filtering, ordering, importing, and exporting, makes it suitable for processing large HD cytometry datasets. While there are R packages which operate on *data.frame* formats (e.g. dplyr [58]), our simple benchmarking measurements show *data. table* to be faster when handling large datasets (Supp. Figure 1). The simple tabular format allows for interoperability between Spectre, basic R functions, and other functions from cytometry or single-cell-specific packages, by storing all the relevant information for each cell in a single, high-performance table. Here, columns desired for use with each function can be easily specified, and new columns that are added as a result of analysis are easily identifiable through the use of regular expression patterns (such as “CD4_asinh”, “CD4_aligned”, etc.). Column metadata in this context is less relevant for processing and analysis, but can still be imported and managed with the main data should the user choose to do so. By designing all our functions to operate on this simple *data.table* structure, we remove the requirements for users to convert their data into specific formats for different functions, greatly improving usability.

### Usage flexibility

We have demonstrated the use of Spectre through various workflows. While they each illustrate an approach to analysis, they are flexible in such a way that functions can be run in any order, and can be replaced with others (including those that are not built into Spectre). For instance, it is possible to run a DR tool on the dataset before clustering, or replacing Spectre’s run kNN classifier function with another function (like a Decision Tree classifier [59]). Additionally, most of Spectre’s functions are designed such that they do not rely on each other to operate. A function may simply be executed on its own without having to run others beforehand. Some functions require data to be in a particular layout with appropriate calculations (e.g. calculating MFI across samples for heatmap creation with make.pheatmap), but this can be achieved either with (using write.sumtables) or without (using other functions or manual creation) Spectre. In addition, Spectre’s functions are equipped with parameters that allow users to customize how they operate. The majority of these parameters come with a set of default values that are based on the original implementation of the tools. The capacity to run Spectre’s functions with either default or custom parameter values make Spectre both simple and customizable.

### Docker for accessibility

Spectre provides wrapper functions to many existing computational tools, and thus require users to install the corresponding R packages before using them. Inadvertently, this creates a cycle of package dependencies as these packages often rely on other existing R packages. Managing such dependencies has proven to be a challenge even to advanced analysts, as there exists a myriad of ways to install, remove, and update packages (e.g. CRAN, Bioconductor [60], devtools [61]). Moreover, some packages rely on users to have other softwares (e.g. Xcode in Mac, Rtools in Windows) or compilers available on their computer. This may pose as a major hurdle for those who are not familiar with R. To address this, we made Spectre available as a Docker image. The image is a prepared environment loaded with RStudio for users to interact with R code (write and run) and all the libraries required by Spectre. By downloading the Docker image and launching it as a self-contained computing environment, users will be able to run their analysis without going through the complicated setup process. While Docker introduces an additional layer between user’s physical computing resources and the analytical tools, previous work by IBM indicates the performance degradation to be negligible [62].

### Versatility of application

The structure of all forms of cytometry data, including flow, spectral or mass, are inherently similar: each row is an individual cell, whilst each column is an individual marker or feature measured on those cells. The values in such a table indicate the signal intensity for each individual marker within the panel on each cell. Thus, Spectre can be used on datasets generated by both flow (including spectral) and mass cytometry, following compensation (or spectral unmixing), and initial cleanup gating. As a result, Spectre can also be used on other forms of cellular data, such single-cell RNA seq data, following some additional pre-processing steps. Additionally, Spectre can be used to analyze HD imaging data, such as that generated by IMC, once cellular segmentation has been performed.

## DATA AVAILABILITY

Spectre code and demonstration data used in this paper are available at https://github.com/immunedynamics/spectre.

## ACKNOWLEDGEMENTS

This work was supported by a grant from the Marie Bashir Institute for Infectious Disease and Biosecurity. This work was supported in part by NH&MRC Project grant 1088242 and grant from the Merridew Foundation to N.J.C.K. Authors T.M.A. and F.M-W. are supported by the International Society for the Advancement of Cytometry (ISAC) Marylou Ingram Scholars program. G.H.P. is supported by the Australian Government Research Training Program (RTP) Scholarship. A.G.S. is supported by the Australian Government RTP Scholarship and The University of Sydney Postgraduate Merit Award. We would like to acknowledge the contribution of our colleagues who inspired the design and implementation of Spectre.

---

**Supplementary Figure 1. Data processing speeds**

A) Benchmarking of files reading/writing with data.table or base R functions. B) Benchmarking of common data operations using data.table or Dplyr. 12,108,595 cells with approx. 25 dimensions were analyzed on a virtual windows machine with 16 GB of RAM.

**Supplementary Figure 2. Data visualization and ArcSinh transformation**

A) Brief comparison of plot visualization options as implemented in FlowJo, with the axis set as linear (with default maximum values), linear (with adjusted maximums), logarithmic, or bi-exponential/logicle.

B) Direct comparison of raw data using a bi-exponential/logicle axis with data subject to ArcSinh transformation and plotted on a linear axis.

**Supplementary Figure 3. ArcSinh transformation of mass cytometry data**

A) Plots of raw data in FlowJo with various width basis (WB) settings for the x-axis (168Er CD8a). Minimum x-axis value for each plot was adjusted to the highest possible value, within the restrictions imposed by FlowJo. Y-axis width basis was fixed at −100. Both x- and y-axis positive decades were set at 4.75. B) Plots of ArcSinh transformed data from R with various co-factors used for the marker on the x-axis. Y-axis co-factor was fixed at 10.

**Supplementary Figure 4. ArcSinh transformation of conventional and spectral cytometry data**

Plots of data generated by a conventional (A-B) and spectral (C-D) flow cytometry system. The conventional flow cytometry system was a 10-laser BD LSR-II, and the spectral flow cytometry system was a 5-laser Cytek Aurora. A) Plots with of conventional flow cytometry raw data in FlowJo with various width basis (WB) settings for the x-axis (BUV805 CD8a). Minimum x-axis value for each plot was adjusted to the highest possible value, within the restrictions imposed by FlowJo. Y-axis width basis was fixed at – 100. Both x- and y-axis positive decades were set at 4.29. B) Plots of ArcSinh transformed data from R with various co-factors used for the marker on the x-axis. Y-axis co-factor was fixed at 1000. C) Plots of spectral cytometry raw data in FlowJo with various width basis (WB) settings for the x-axis (BUV 805 CD8a). Minimum x-axis value for each plot was adjusted to the highest possible value, within the restrictions imposed by FlowJo. Y-axis width basis was fixed at –631. Both x- and y-axis positive decades were set at 5.68. D) Plots of ArcSinh transformed data from R with various co-factors used for the marker on the x-axis. Y-axis co-factor was fixed at 5000.

**Supplementary Figure 5. ArcSinh, normalized, and noise reduced data**

Transformations and manipulations performed on conventional flow cytometry data. A) Raw data plot of FITC Ly6C vs BUV805 CD8a. B) ArcSinh transformed data (a-axis) using a co-factor (CF) of 1000. C) ArcSinh transformed data (x-axis) redistributed between 0 and 1 using the *do.normalise* function. D) ArcSinh transformed data (a-axis) with data points below 0 changed to 0 using the *do. noise. reduce* function, to reduce the distribution of negative cells below zero.

**Supplementary Figure 6. Dimensionality reduction crowding and computation time**

A-B) Plots of tSNE analysis performed with different numbers of total cells, colored by relative density, or cell type. C) Computation time for tSNE on different sized datasets, plotted on a logarithmic scale.

**Supplementary Figure 7. Evaluating kNN classification accuracy**

A) A single labelled dataset was split into halves – one half retaining population labels (training data) and one half having population labels hidden (validation data). B) A kNN model was trained on the training data and applied to the validation data. C) Comparison of the original labels on the training data and the predicted labels on the validation data. D) A graph showing the accuracy for the kNN classifier for different values of *k*.

