## Supplementary Figures for "Integration, exploration, and analysis of high-dimensional single-cell cytometry data using Spectre"

#### Supplementary Figure 1

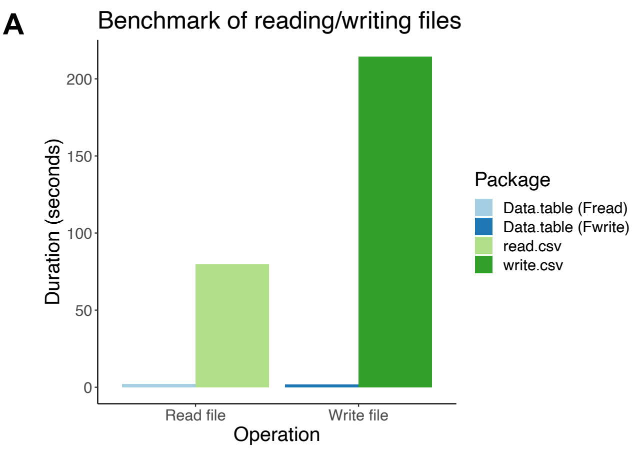


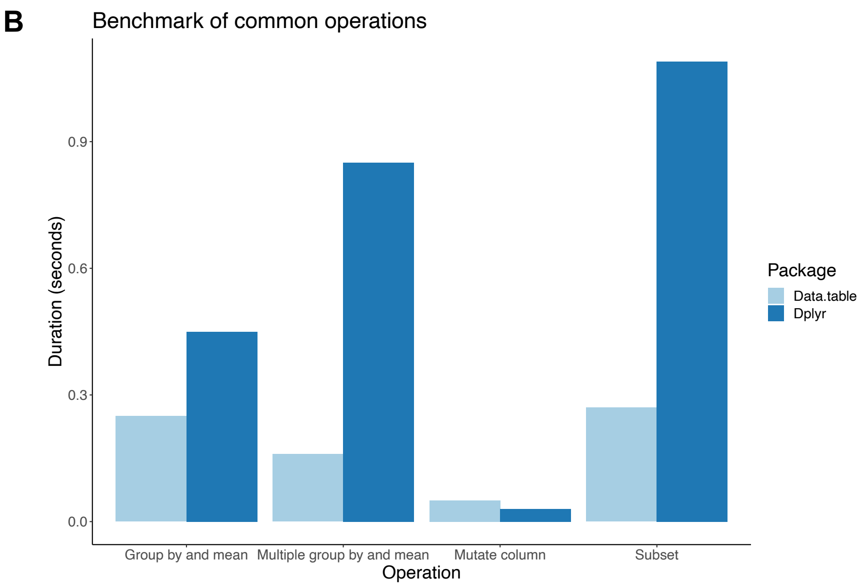

### Supplementary Figure 2

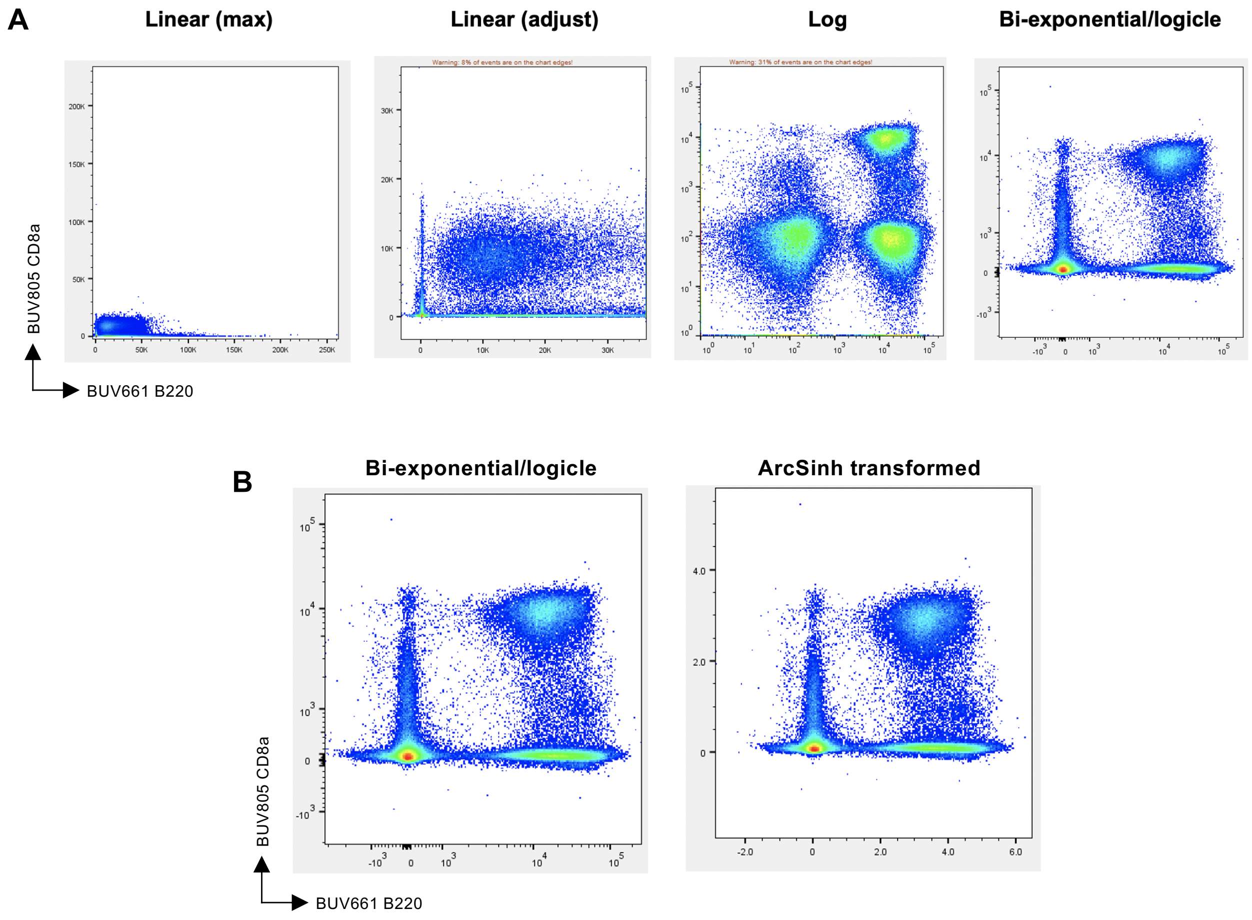

#### Supplementary Figure 3

**A** Varying width basis for biexponential plotting of x-axis (CD8a)

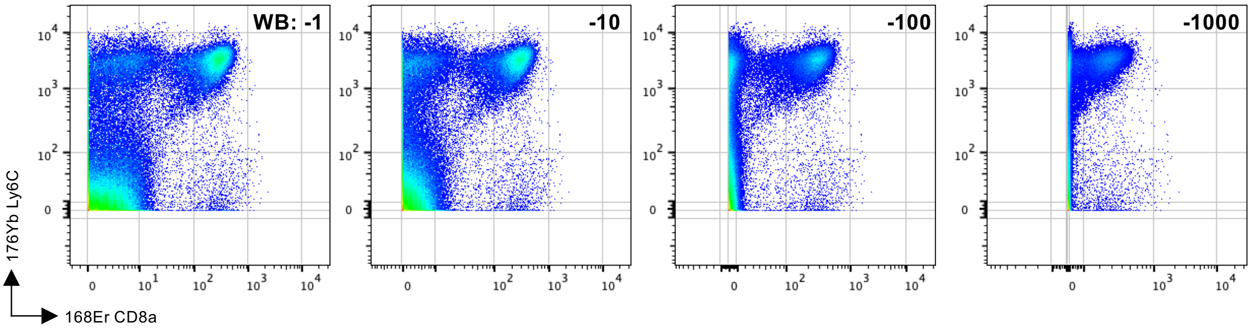


**B** Varying co-factor for ArcSinh transformation of x-axis (CD8a)

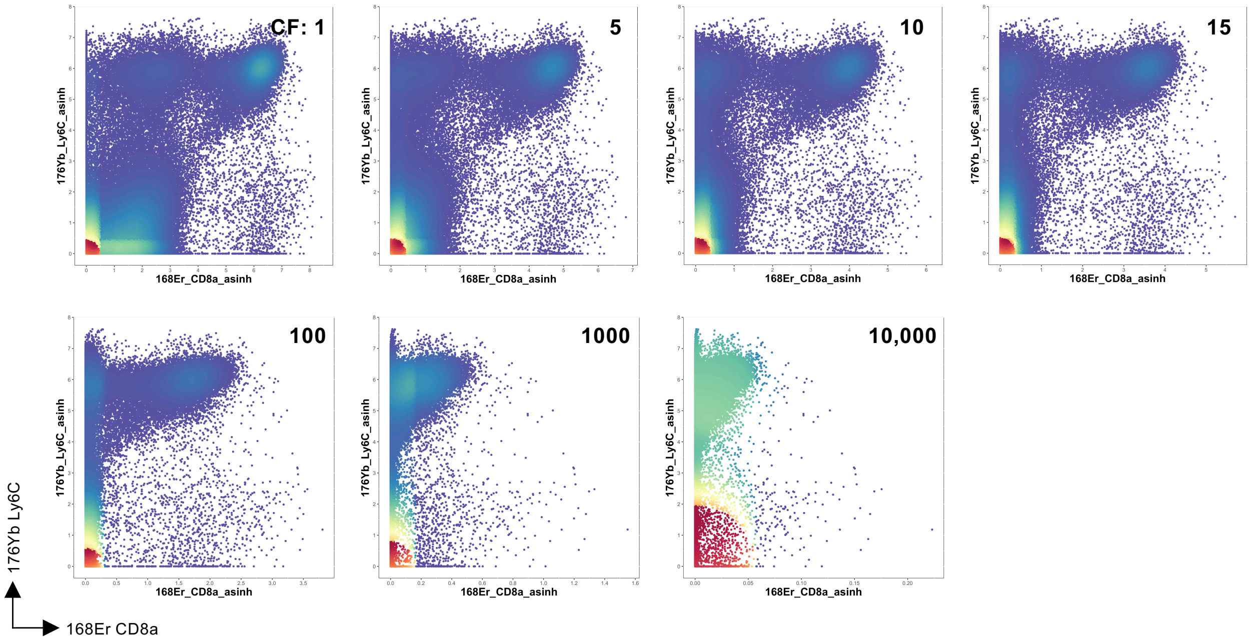

#### Supplementary Figure 4

##### Conventional flow cytometer (10-laser BD LSR-II)

**A** Varying width basis for biexponential plotting of x-axis (CD8a)

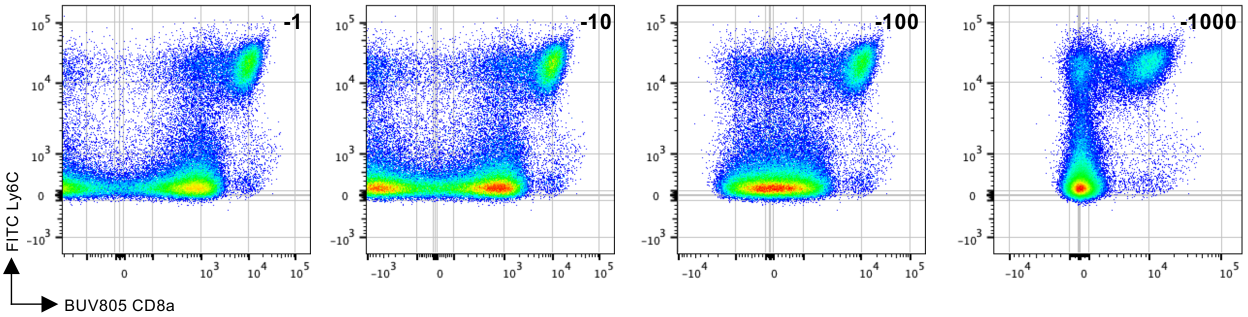


**B** Varying co-factor for ArcSinh transformation of x-axis (CD8a)

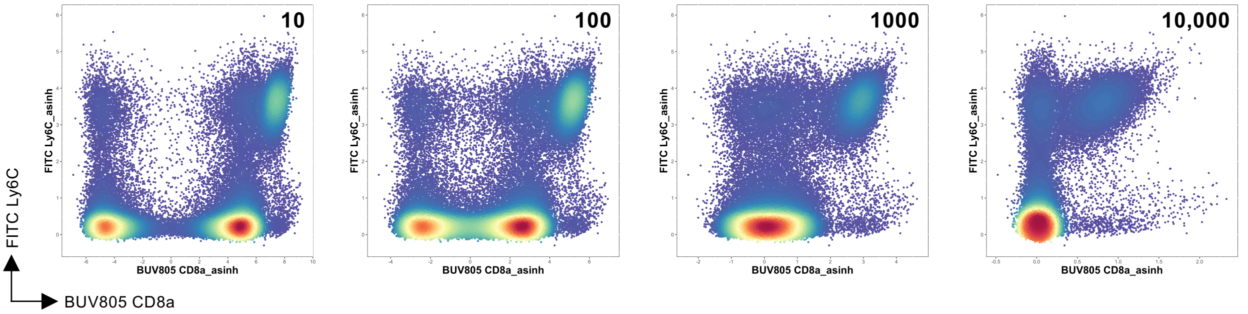


##### Spectral flow cytometer (5-laser Cytek Aurora)

**C** Varying width basis for biexponential plotting of x-axis (CD8a)

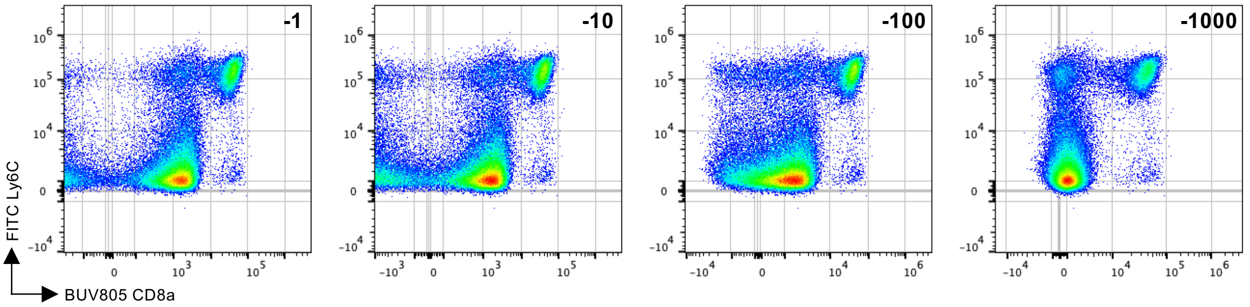


**D** Varying co-factor for ArcSinh transformation of x-axis (CD8a)

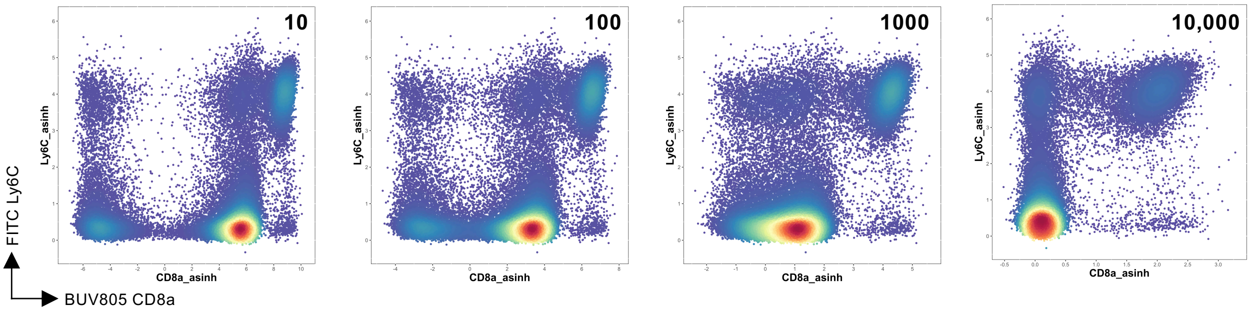

### Supplementary Figure 5

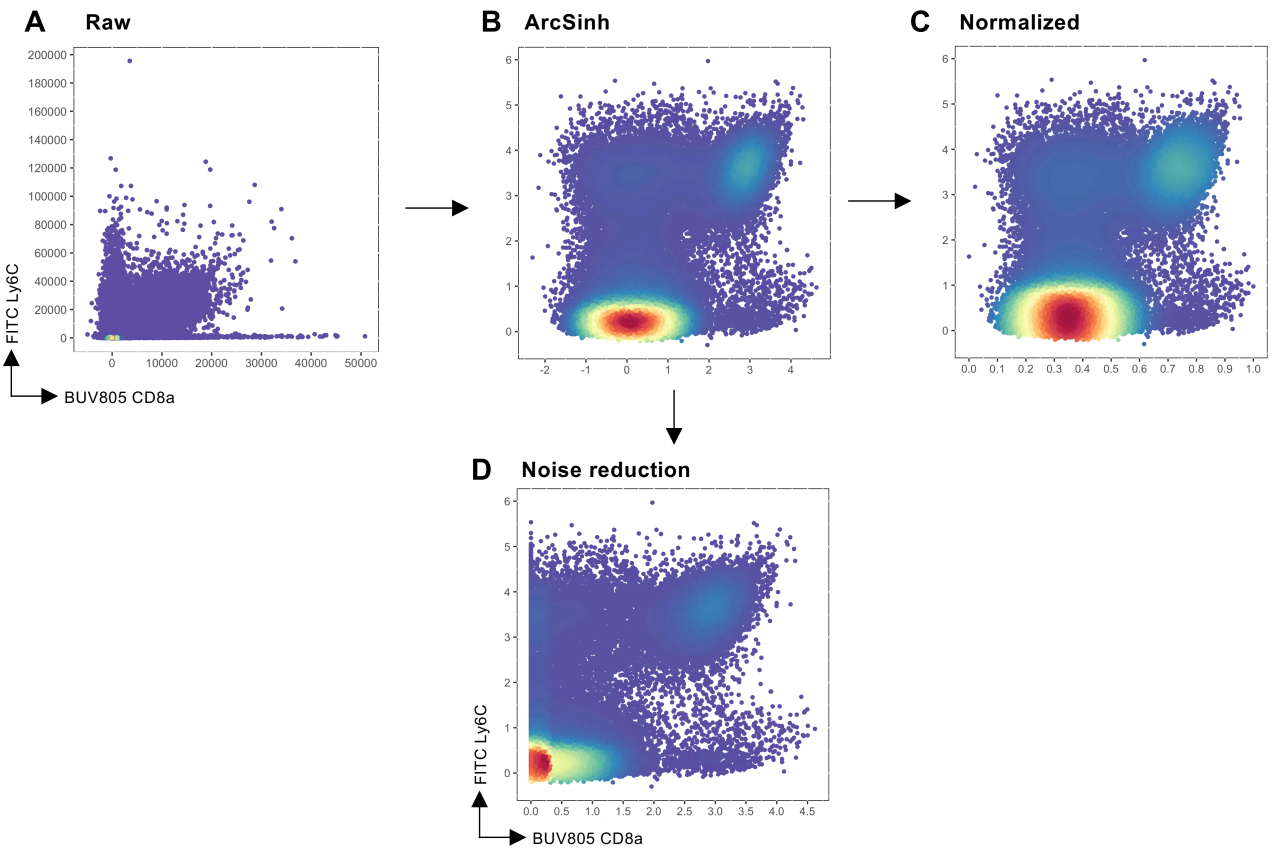

#### Supplementary Figure 6

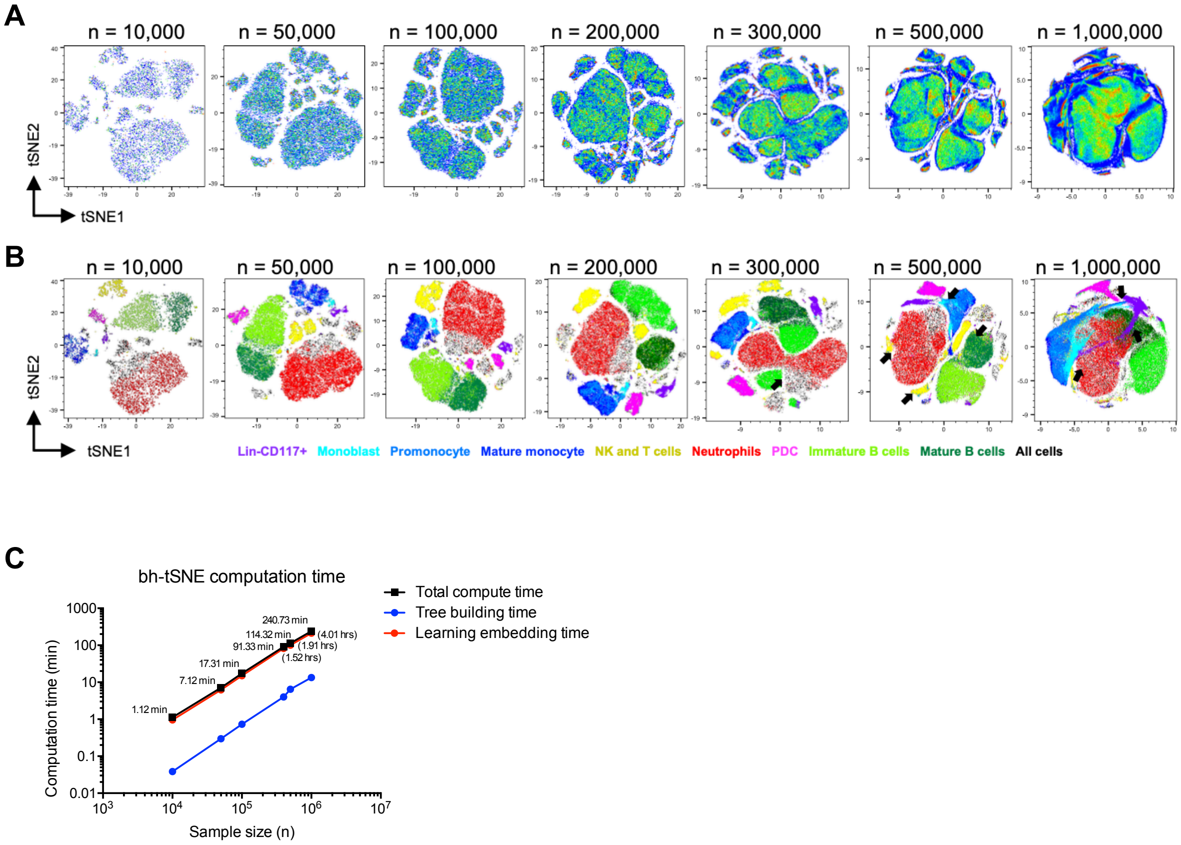

#### Supplementary Figure 7

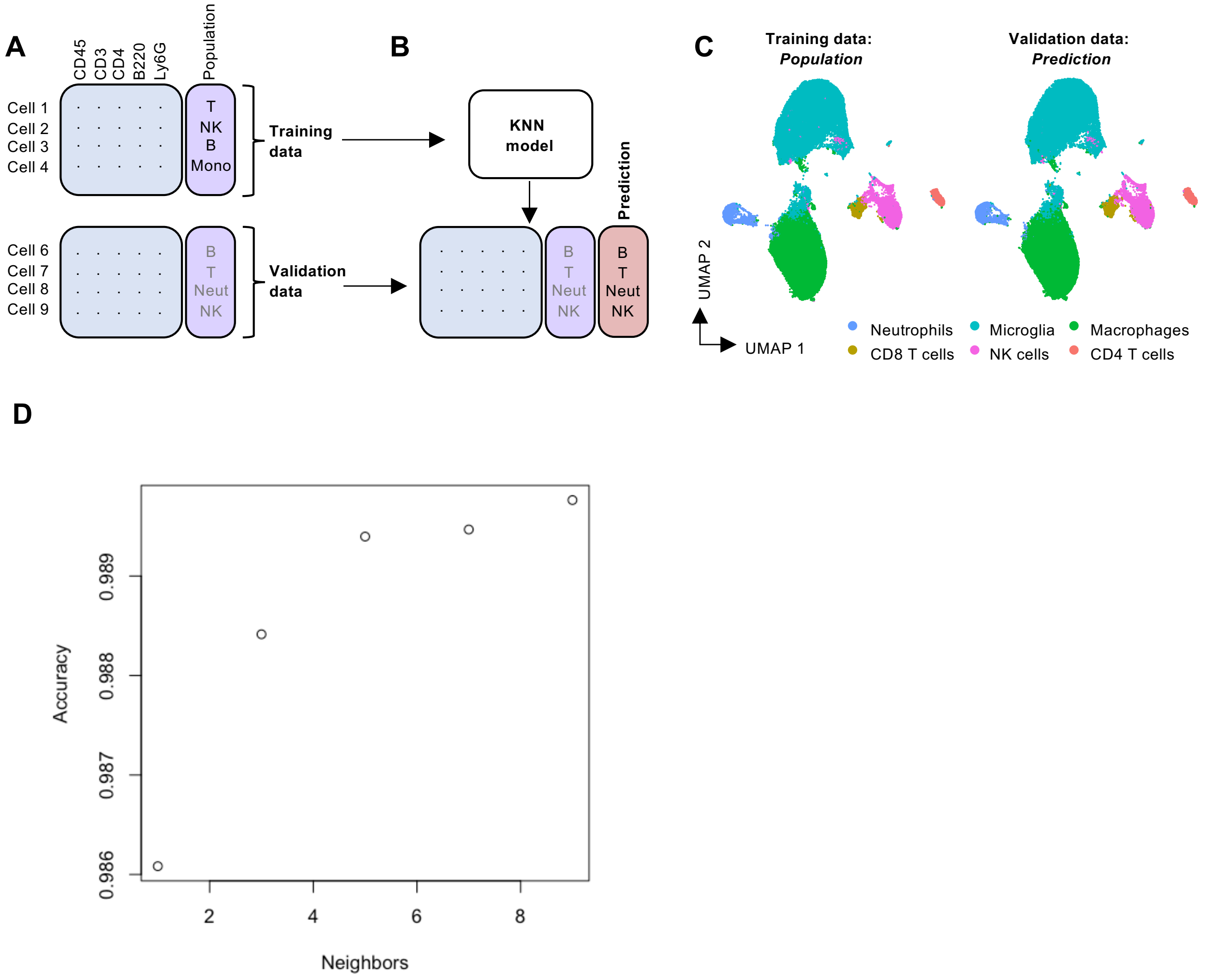
